# Comparison of BOLD and CBV using 3D EPI and 3D GRASE for cortical layer fMRI at 7T

**DOI:** 10.1101/778142

**Authors:** Alexander JS Beckett, Tetiana Dadakova, Jennifer Townsend, Laurentius Huber, Suhyung Park, David A Feinberg

**Author notes:** Corresponding Author: Alexander Beckett, Helen Wills Neuroscience Institute, University of California, Berkeley Berkeley, CA.

## Abstract

**Purpose:** Functional MRI (fMRI) at the mesoscale of cortical layers and columns requires both sensitivity and specificity, which can be compromised if the imaging method is affected by vascular artifacts, particularly cortical draining veins at the pial surface. Recent studies have shown that cerebral blood volume (CBV) imaging is more specific to the actual laminar locus of neural activity than BOLD imaging when using standard gradient-echo (GE) EPI sequences. Gradient and Spin Echo (GRASE) BOLD imaging has also shown greater specificity when compared with GE-BOLD.

**Methods:** Here we directly compare CBV and BOLD contrasts in high-resolution imaging of the primary motor cortex for laminar fMRI in four combinations of signal labeling, VASO (CBV) and BOLD with 3D GE-EPI and zoomed 3D GRASE image readouts.

**Results:** We find that both CBV imaging using EPI-VASO and BOLD imaging using GRASE-BOLD, show similar specificity and sensitivity and are thus useful tools for mesoscopic fMRI in the human cortex.

**Conclusion:** These techniques demonstrate sufficient sensitivity and specificity to allow layer-fMRI to be used by neuroscientists in a wide range of investigations of depth-dependent neural circuitry in the human brain.

## Introduction

High-resolution imaging of neural architecture at the mesoscale is growing in importance, especially due to cortical laminar analysis for understanding neural circuits. Blood oxygen level-dependent (BOLD) contrast is the most commonly used functional magnetic resonance imaging (fMRI) method to investigate neural activity in the human brain (1). Recent results using Gradient And Spin Echo (GRASE) imaging (2, 3), where orthogonal excitation and refocusing pulses are used to image a smaller area of cortex with higher resolution and specificity, have demonstrated the ability to resolve neuronal organization at the mesoscale (4–6). Alongside this, Cerebral Blood Volume (CBV) imaging using the VAscular Space Occupancy (VASO) technique (7, 8) have demonstrated the ability to resolve activity from different cortical layers (9, 10). In this paper, different combinations of image readout (EPI/GRASE) and contrast (BOLD/CBV) are compared for sensitivity and specificity in laminar imaging.

BOLD contrast is a combination of blood T2* or T2 changes and signal dephasing due to susceptibility changes between blood and surrounding tissues following neural activation (1). The BOLD signal arises from both small vessels specific to local neural activity, and draining veins that are not locally specific (11). In both small and large vessels, the BOLD signal consists of multiple components: intravascular (IV), arising from T2 and T2* effects of blood oxygenation induced susceptibility effects, and extravascular (EV), arising from static dephasing induced by field inhomogeneities and dynamic averaging around small vessels. Whilst the indirect nature of the BOLD response, being based as it is on changes in hemodynamics (12), prevents measuring the activity of individual neurons, it has the potential of measuring the activity of cohorts of neurons arranged at the mesoscale (0.5 – 1mm), namely neurons arranged radially (columns) and tangentially (layers) to the cortical surface. Cortical columns are groups of neurons arranged perpendicularly to the cortical surface that share a common preference for a stimulus (13, 14), for example ocular dominance columns in V1 (15, 16). In addition to this radial organization, the cortical sheet is divided parallel to the cortical surface into roughly six layers, based on the properties and connections of the cells contained within each layer. The connections between these layers within a patch of cortex, and the connections of these layers to other brain regions, are thought to define the processing performed in that region of cortex, with bottom-up and top-down information flow being defined by its laminar target (17). Being able to non-invasively measure neural activity at this scale would allow important insights into the fundamental circuits underlying cognition.

Increases in field strength and advances in hardware now allow sufficient signal to image the brain at mesoscale resolution, and at this level it is the origins of the BOLD signal itself that define the sampling resolution for fMRI. When performing fMRI at ultrahigh field (UHF, 7T and higher), the signal change from the EV component increases quadratically with field strength (1,18,19), while the IV signal in both small and large vessels significantly decreases because the T2* or T2 of blood is very short (18, 20), leading to some increase in specificity. However, with the most commonly used gradient echo (GE) echo planar imaging (EPI) acquisitions at ultrahigh fields, the EV BOLD contrast from large vessels will still be present in the images. This is seen most commonly as bias towards the outer (pial) surface of the neocortex when measuring signal change at different cortical depths (21, 22). Whilst careful experiment design, analysis and post-processing can recover some of the required specificity from GE BOLD acquisitions for laminar (22, 23) or columnar (15,16,24,25) imaging, the specificity of the GE signal has not allowed robust mesoscale mapping to reveal neuronal circuitry in cortical columnar or laminar organization.

Using Spin Echo (SE) EPI for fMRI, a 180° refocusing pulse partially recovers signal loss from static field dephasing in the EV signal around large vessels (19,20,26). This reduces the venous contribution to BOLD, which increases the spatial specificity (19, 20), but at the expense of overall lower sensitivity. However, combining SE imaging with a high B_0_ results in suppression of both the EV and IV components in large vessels (19,20,27), leading to further increases in specificity. SE EPI acquisitions have been used successfully for fMRI at 7T for imaging layers in animals (12,28,29) and columns in humans (16, 30). When using an EPI readout for SE imaging, high-resolution imaging with an extended field of view (FOV) in the phase encoding direction requires very long readout times, which introduces increasing T2* weighting at the outer regions of k-space, reducing image specificity. Segmented (16, 30) or accelerated (4) imaging can shorten echo trains to mitigate this effect, at the expense of increased noise or reduced signal. An alternative and complimentary approach is inner volume (zoomed) imaging, where a smaller FOV (31) is acquired to limit the length of the overall echo train. This is achieved in the SE EPI pulse sequence by orthogonalizing the excitation and refocusing pulse planes, reducing FOV to a limited area of cortex (16, 30).

GRASE (2, 3) combines multiple SE refocusing pulses of a Carr-Purcell-Meiboom-Gill (CPMG) sequence with intervening EPI echo trains in single shot imaging. Sub-millimeter resolution fMRI can be achieved by combining inner volume zooming with single-shot 3D GRASE (2) to reduce the FOV and shorten the multiple EPI echo trains to minimize the T2* weighting (4). Zoomed 3D-GRASE has been used for both laminar (5,21,32,33) and columnar specific imaging (6,32,34). The multiple short echo trains of GRASE increase T2 weighting and decrease T2* weighting in outer k-space. Another difference from SE-EPI is that GRASE is a variant of the CPMG sequence and therefore can contribute stimulated echoes (STE) with T1 contrast to BOLD mechanisms.

Cerebral blood volume (CBV) contrast in fMRI has been shown to have an advantage over BOLD by excluding venous signal contributions and has been shown to be tightly coupled to metabolic changes associated with neural activity (35–37). Non-invasive CBV-weighted fMRI in humans can be achieved using the Vascular Space Occupancy (VASO) method (7, 8), which measures changes in CBV by acquiring an image around the short period while blood signal is nulled after an inversion pulse, so that changes in blood volume lead to a proportional decrease in signal. Recently the technique has been used successfully at UHF (38) for laminar specific applications (9, 10). 3D GRASE can also be used as readout sequence in CBV imaging, as used at low resolutions for fMRI at 3T (39–41) and in resting state experiments (42, 43). However, VASO 3D GRASE has not been developed or evaluated for mesoscale high resolution CBV-fMRI.

In this work, we sought to compare CBV-fMRI using VASO with both 3D GE-EPI and zoomed 3D GRASE readouts for high-resolution layer-dependent fMRI. For both VASO sequences, a double image sequence was utilized as previously for 7T laminar VASO imaging (7,9,10). This double image acquisition gives rise to two FMRI time-series data sets acquired simultaneously: a blood nulled VASO time-series and a standard BOLD time-series. The second image is used to remove the BOLD contrast that exists in the first (blood-nulled VASO) image, a particular issue at high-field for both sequences. We hypothesize that the T2-weighted 3D GRASE BOLD images would show layer profiles similar to the VASO images, with both having less pial surface bias than T2*-weighted 3D EPI BOLD.

The terms used for the four different image/contrast types throughout the manuscript are summarized in Table 1.

**Table 1).**
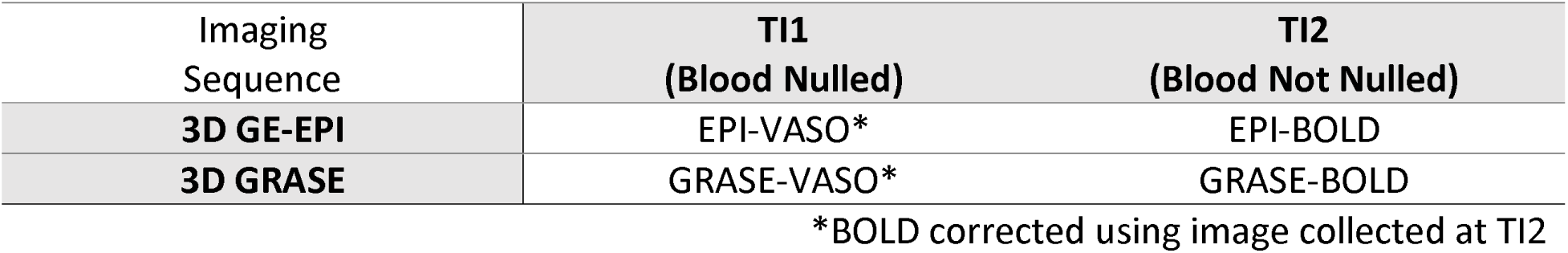
Simplifying terms used for the four different image/contrast types.

## Methods

### Data Acquisition

We analyzed activation-related signal change in the region of motor cortex representing finger and thumb motion, and activation-related signal changes as a functional of cortical depth, for all four combinations of contrast and acquisition (i.e., EPI-BOLD, EPI-VASO, GRASE-BOLD and GRASE-VASO).

#### Imaging Hardware

FMRI data was acquired on a MAGNETOM 7T scanner (Siemens Healthineers, Erlangen, Germany), with an SC72 body gradient coil (Gmax = 70mT/m and SR= 200 mT/m/ms, effectively set to Gmax = 42mT/m and SR= 190 mT/m/ms by inbuilt scanner software limits). RF reception and transmission were performed with a 1 Channel Transmit/32 Channel Receive Head Coil (Nova Medical, Wilmington, MA, USA). The study protocol was approved by the Institutional Review Board at the San Francisco VA Center; each participant gave written informed consent before MRI data acquisition. 6 healthy volunteers (age: 43.2 ± 13.7, 3 female) participated in this study.

#### Stimulation paradigm

To induce VASO and BOLD functional signal changes, a unilateral finger tapping task (thumb and index finger) paradigm, previously used to investigate layer specific activation, was utilized (10). In brief, it consisted of 12 blocks, each of 60 s duration (30 s tapping, 30 s resting), resulting in acquisition time (TA) of 12 min. Subjects were cued to tap their forefinger and thumb with the same pacing as a video animation (44) projected in the scanner bore. This paradigm leads to an increase in activity in Layer II/III (cortical input) and Layer Va (spinal output) in the hand-knob region in M1 (10), and this expected pattern of activation can be used to assess the specificity of the different sequences used.

#### Pulse Sequence

A slice-selective slab-inversion (SS-SI) VASO sequence (7) with GRASE (3) readout was implemented. Specifically, the GRASE pulse sequence was modified to include an inversion recovery (IR) pulse to acquire an image with VASO contrast, followed by an additional readout to acquire a second image with BOLD contrast (Fig. 1). SS-SI VASO EPI was implemented as per previous studies (10, 45) with a 3D GE-EPI readout (46). In VASO, the signal decreases with an increase in blood volume, so CBV change is inversely proportional to the VASO signal change, and is expressed as CBV change in ml per 100ml of tissue (9, 47). In the current experiment results were expressed in units of percent signal change to allow easier comparison with the BOLD results and across multiple sequences.

**Figure 1.**
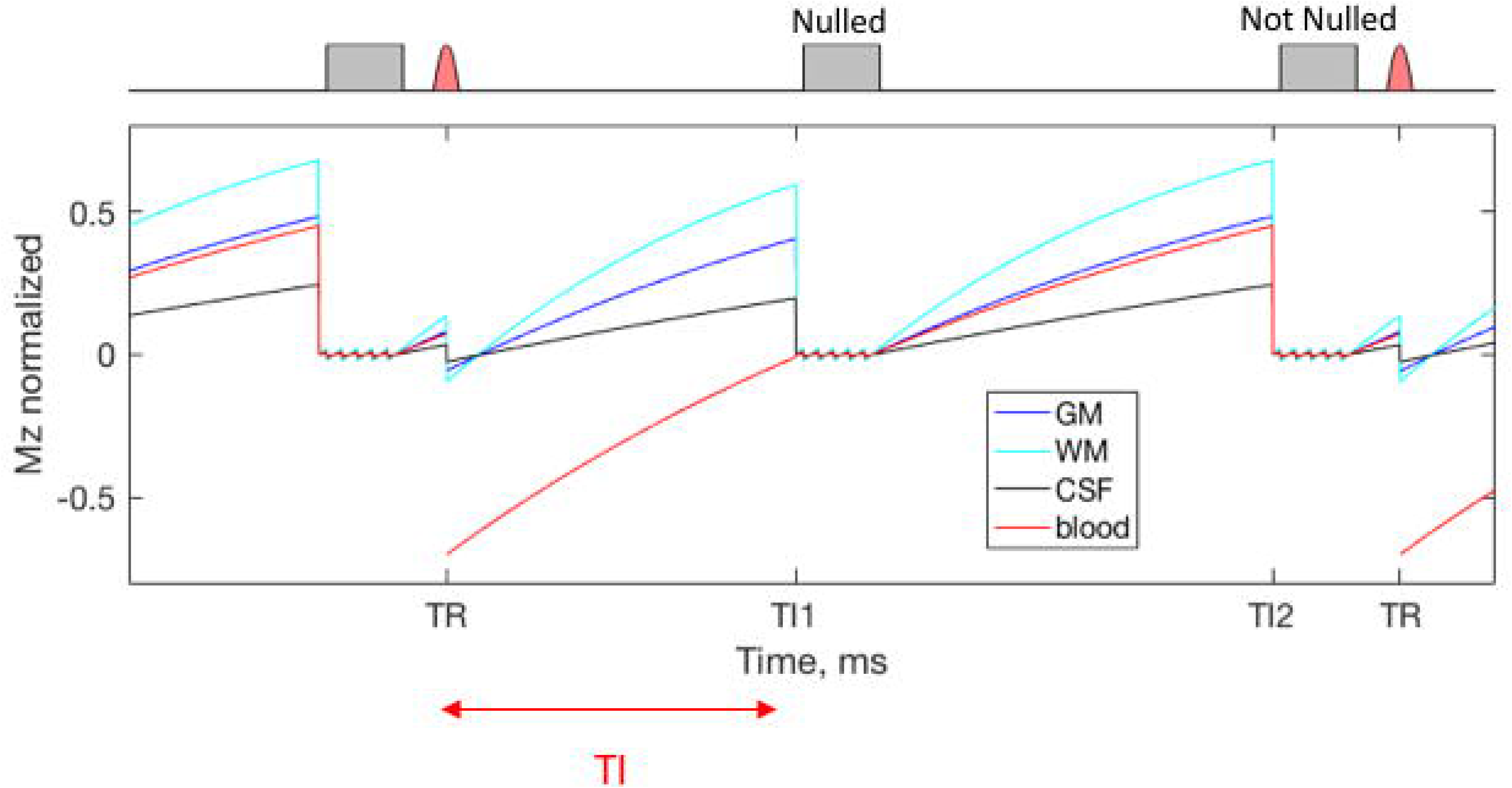
GRASE pulse sequence with two readouts (one with blood nulled, one without nulling), which are used to provide VASO and BOLD functional contrast. Blood signal (red curve) is assumed to not be in steady-state. The signals of grey matter, white matter, and cerebrospinal fluid (blue, turquoise and black curves) are assumed to be in steady state after previous inversions and readouts, whereas fresh blood is inverted every TR. Inversion recovery pulses are drawn with red and GRASE image readouts are drawn in gray. At the time of the Nulled image acquisition, blood signal is nulled, whereas at Not Nulled image acquisition, there is signal for all tissue types.

Comparison between the 3D GE-EPI (referred to simply as EPI below) and 3D GRASE (referred to as GRASE below) readouts is shown in Figure 2. In EPI, the 3D volume is excited using a small flip angle (FA) for each phase encoding step along the slice (partition) direction. The signal decays with T2* along the in-plane phase encoding direction, so the final 3D-image has T2* contrast. The signal acquisition ordering in the partition direction was linear (center-out-center), with k_0_ acquired at the time after the inversion pulse (inversion time, TI) at which blood signal was nulled, and the outer k-space partitions collected slightly before or after the blood nulling TI. In previous comparisons of linear and centric k-space ordering with k0 at the blood nulling TI, linear ordering was found to be more robust, hence was used here. In GRASE, the 3D volume is excited only once and then the magnetization is refocused at each partition using 0 nominal 180 RF pulses, with phase encoding in the partition direction. Along the partition direction the signal decays with T2, and spin echoes are refocused at the k-space center for each partition. Between refocused echoes (corresponding to the outer k-space phase encoding lines) the signal undergoes T2* decay as well. Images acquired with GRASE readout have combined T2* and T2 contrast, improving specificity.

**Figure 2:**
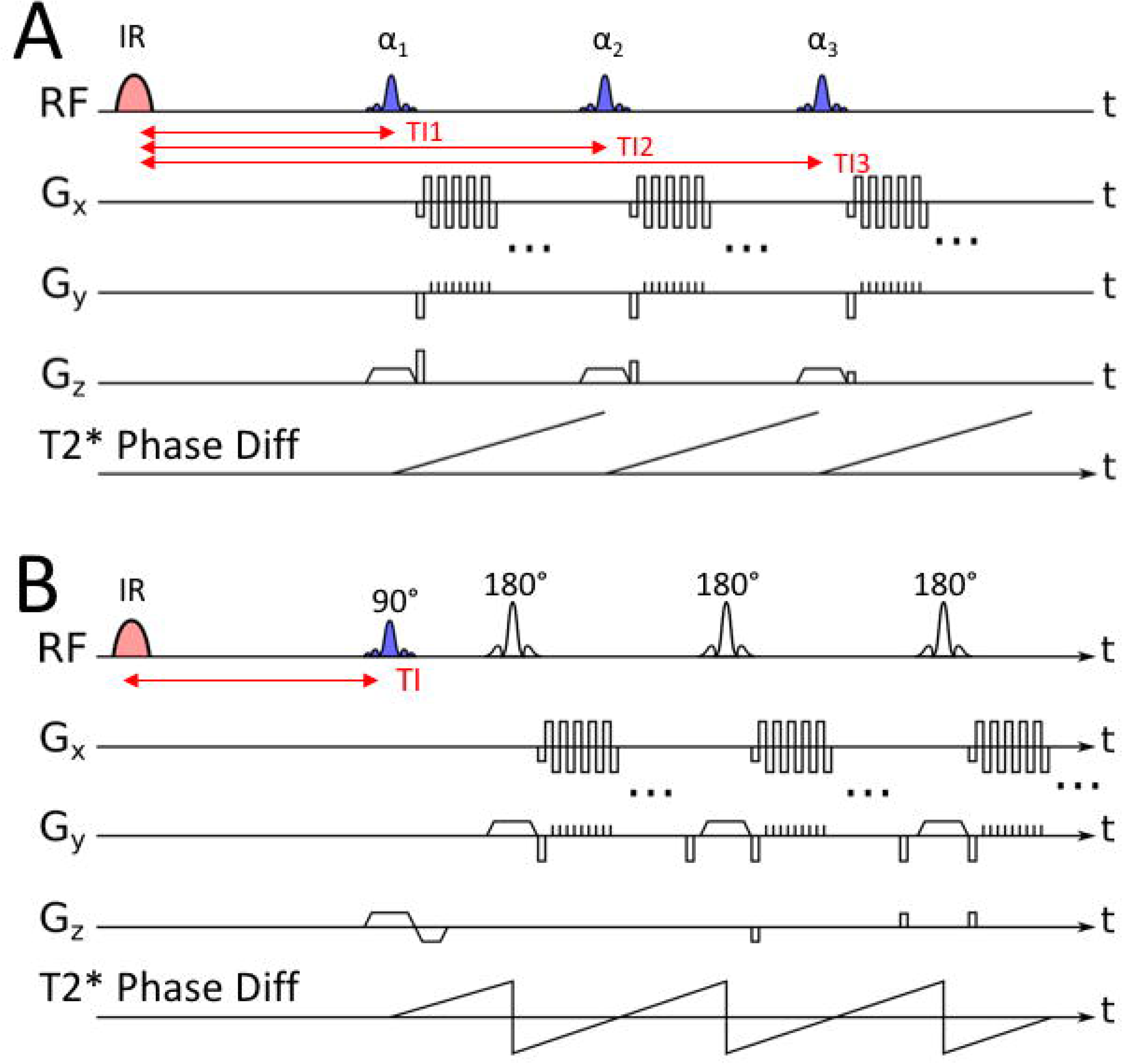
Comparison of two acquisition methods. A: 3D gradient echo EPI, B: 3D GRASE. Note that 3D EPI image acquisition involves multiple excitation pulses at slightly different TIs, which can lead to variable T1 weighting across the k-space partitions. 3D GRASE has only a single excitation, leading to uniform T1 weighting across k-space. GRASE greatly reduces T2* in the image compared to EPI, given signal dephasing differences (T2* Phase diff) are refocused multiple times to zero phase error, where spin echoes are created.

With the IR pulse necessary for VASO contrast, the EPI acquisition will have each k-space partition acquired at progressively later TIs, leading to different T1 weightings across the slice-partition direction, which can result in blurring, and is mitigated by using variable FA (VFA) across the partition direction (48), assuming gray matter (GM) T1 of 1800ms. This VFA method can only correct for the blurring in a single tissue component (GM) whereas white matter (WM) and Cerebral Spinal Fluid (CSF) have different T1s. In addition, the VFA approach is limited by the inhomogeneities in RF transmit field at 7 T, hence, the T1-related blurring effect might be only partially corrected. GRASE signal, on the other hand, will be acquired at a single TI, established in time by the single excitation pulse, throughout the whole 3D k-space, so all partitions in the 3D volume will have exactly the same T1 weighting. While blood signal will remain nulled across the echo train, signal in other tissues will be affected by T2 decay across the echo train, causing signal variations across slice partitions and hence blurring in the slice direction (4), which can be mitigated to an extent by the use of VFA for the refocusing pulses at the cost of SNR (49). Excitation FA for the GRASE readout was set to 90°, while the refocusing FA had to be decreased to 165° to reduce power deposition and specific absorption rate (SAR) given the required 1.5s between image readouts. VFA would have reduced the PSF blurring in 3D GRASE compared to using constant refocusing pulses (49), and was not used in these initial experiments to simplify comparisons. VFA refocusing is prone to B1 inhomogeneity signal losses (49), and will introduce signal from STE into the image. The constant refocusing flip angles in 3D GRASE only hold if there is a uniform B1 field in the imaging region, and any imperfections in these nominal flip angles will have a deleterious effect on blurring for both sequences.

To minimize the duration of the EPI readout and therefore the T2* decay, while achieving high spatial resolution, an inner volume (zoomed) acquisition (31) was implemented for GRASE (Fig. 3 The FOV for GRASE was 99×25×12 mm^3^ and matrix size was 132×34×8, yielding a nominal 3 resolution of 0.75×0.75×1.5 mm^3^; TE was 48ms. Partial Fourier sampling of 5/8 was used in partition direction to reduce the total echo train length to minimize T2 blurring. To minimize TE and maximize SNR for the center of k-space, centric reordering was implemented in the 3 partition direction. EPI had FOV 98×32.8×12 mm^3^, with a matrix size of 132×44×8, for a nominal 3 resolution of 0.75×0.75×1.5 mm^3^; TE was 24ms. Partial Fourier of 6/8 and GRAPPA acceleration of 2 were used in the in-plane phase encode direction. Note that while GRASE and EPI have similar FOVs and slice placement, read and phase encoding directions (Gr and Gp) were orthogonal for the two sequences to avoid aliasing along the phase encode direction in EPI. Although this leads to differing phase encode FOV and direction for the two sequences, both are optimal given the requirements and limitations for the two sequences – EPI requires a large phase encode FOV (with acceleration) to avoid wrap-around aliasing of the signal, while GRASE can use zooming to limit the phase encode FOV, minimizing T2* contamination without the need for acceleration and the concomitant SNR penalty.

**Figure 3:**
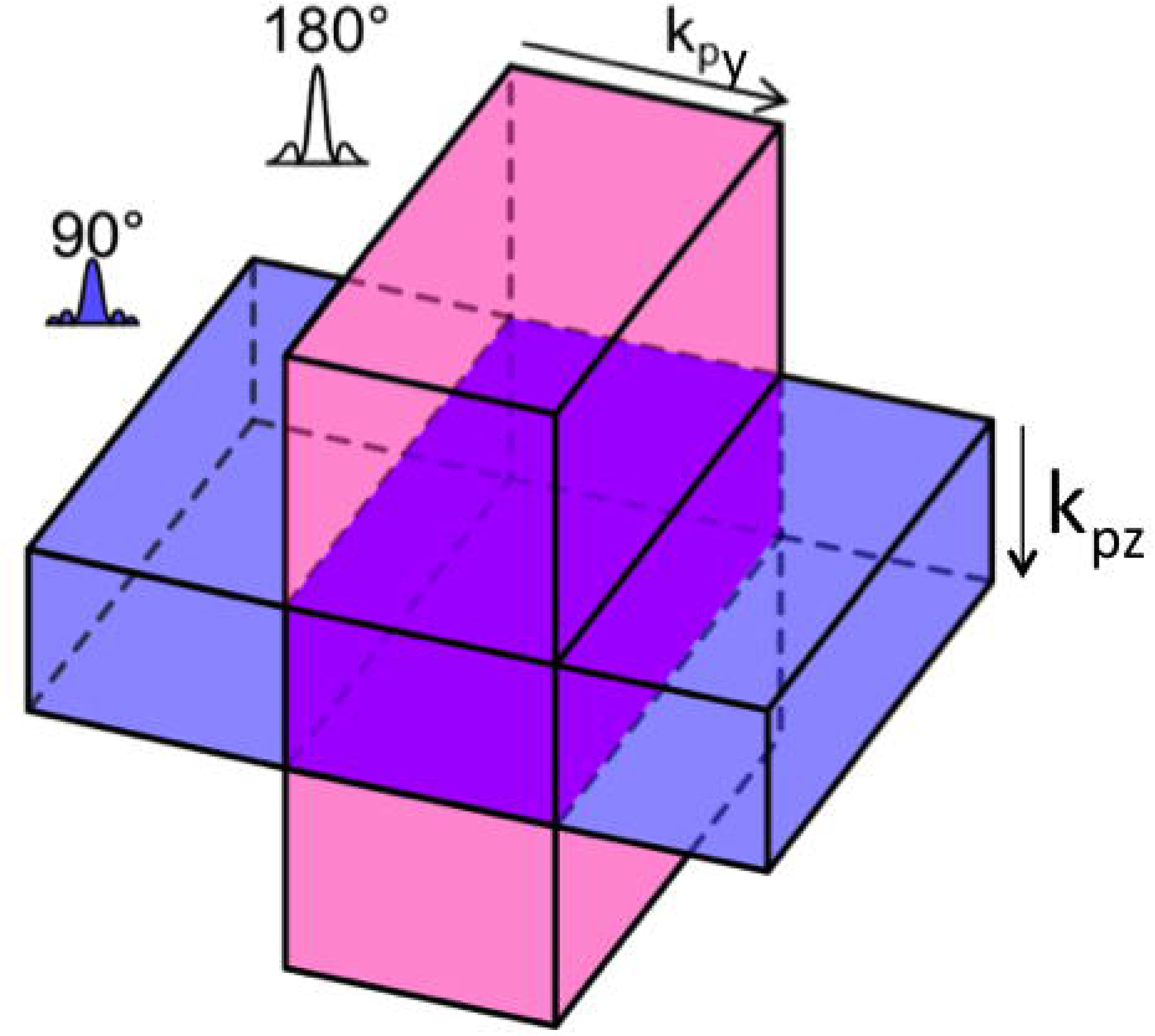
Schematic of inner volume selection using 90° and 180° RF pulses, applied perpendicularly to each other. This limits the FOV in the inplane (y) and partition (z) directions, allowing high resolution acquisition with minimal echo train length, minimizing T2* weighting in the image.

At 7T the blood nulling TI is 1457ms (assuming blood T1 = 2100ms (50)), which is very similar to blood arrival time in the motor cortex (while differences between arterial and venous blood T1 have been reported, these small differences are unlikely to affect the VASO contrast so we assume a single blood T1 (9)). To avoid the non-nulled blood arriving to the motor cortex during VASO image acquisition, an IR pulse with lower inversion efficiency was used for the GRASE acquisition. Specifically, a BIR4 pulse (51) was adapted to have a 71% efficiency (FA = 135°), which decreased the blood nulling TI to 1128 ms. We used TI = 1100 ms in these experiments, as slight deviations from blood nulling TI were shown to not substantially affect the VASO contrast (9). The following timing parameters were used: TI/TR = 1100/3000 ms, Acquisition Time (TA) = 12 min. Time between all imaging readouts was kept constant and was TR/2, as in previous VASO implementations to allow BOLD correction using interleaved Blood Nulled and BOLD acquisitions (9, 10). Fat Suppression was implemented for both sequences in the form of a gaussian saturation pulse (with spoiler) before each excitation pulse.

Table 1 shows the imaging parameters for both sequences.

To increase SNR, the relatively predictable folding pattern of the “hand-knob” gyral pattern in M1 allowed thicker slices to be used when the slices were perpendicular to the central sulcus. Slice position was adjusted for each subject to be perpendicular to the thumb/forefinger region of their M1 hand-knob, based on a separate (0.8mm isotropic voxel size) MP2RAGE image. To mitigate B1+ inhomogeneities that can negatively affect the image quality of spin-echo sequences, and to help ensure a proper inversion pulse for blood nulling, a passive B_1_^+^ shimming approach was adopted by placing high permittivity dielectric pads around the head and neck (52). Transmitter reference voltage was adjusted for each subject by measuring the mean B_1_^+^ values in the volume of interest with the dielectric pads applied, using a B1 mapping procedure.

### Analysis

#### Pre-processing

Image volumes with VASO and BOLD contrast were separately corrected for motion using SPM12 (Functional Imaging Laboratory, University College London, UK), with the option of spatial weighting to optimize correction over the motor cortex. A 4th order spline was used for motion estimation and resampling to minimize blurring (10).

At high field strengths, the positive BOLD signal outweighs the negative VASO signal caused by an increase in CBV (53), for both EPI and GRASE imaging. Dynamic division of the VASO and BOLD volumes was performed to remove residual BOLD contrast contamination from the VASO images, under the assumption that T2* contrast is the same in images with both contrasts because they were acquired concomitantly (7).

#### Functional and layer analysis

The functional activations were calculated as the difference between mean signal during the task and mean signal during rest, ignoring the initial 9s of each period to minimize the influence of transition periods. This method has been shown to provide results that are easier to interpret than methods using inferential statistics, which can be affected by laminar differences in noise and hemodynamic response function (HRF) shape (10).

Functional data were used to create the WM and CSF borders for the laminar analysis to avoid issues of distortion, registration and interpolation (54). To create masks of cortical depths, images were 5 times upsampled (nearest neighbor interpolation) and the GM boundaries with WM and CSF were manually delineated on the anterior bank of the central sulcus, including the hand-knob region of M1 (Fig 4A). Functional VASO images can be used to generate a T1 weighted anatomical image that provides good contrast between GM and WM, which were used as anatomical references for grey matter boundaries in the EPI images (10). For GRASE scans the mean BOLD images showed greater tissue contrast, so these images were used for boundary identification. This delineation in functional space removes the need for distortion correction and co-registration, avoiding registration errors and resolution degradation (blurring) due to resampling (54).

**Figure 4:**
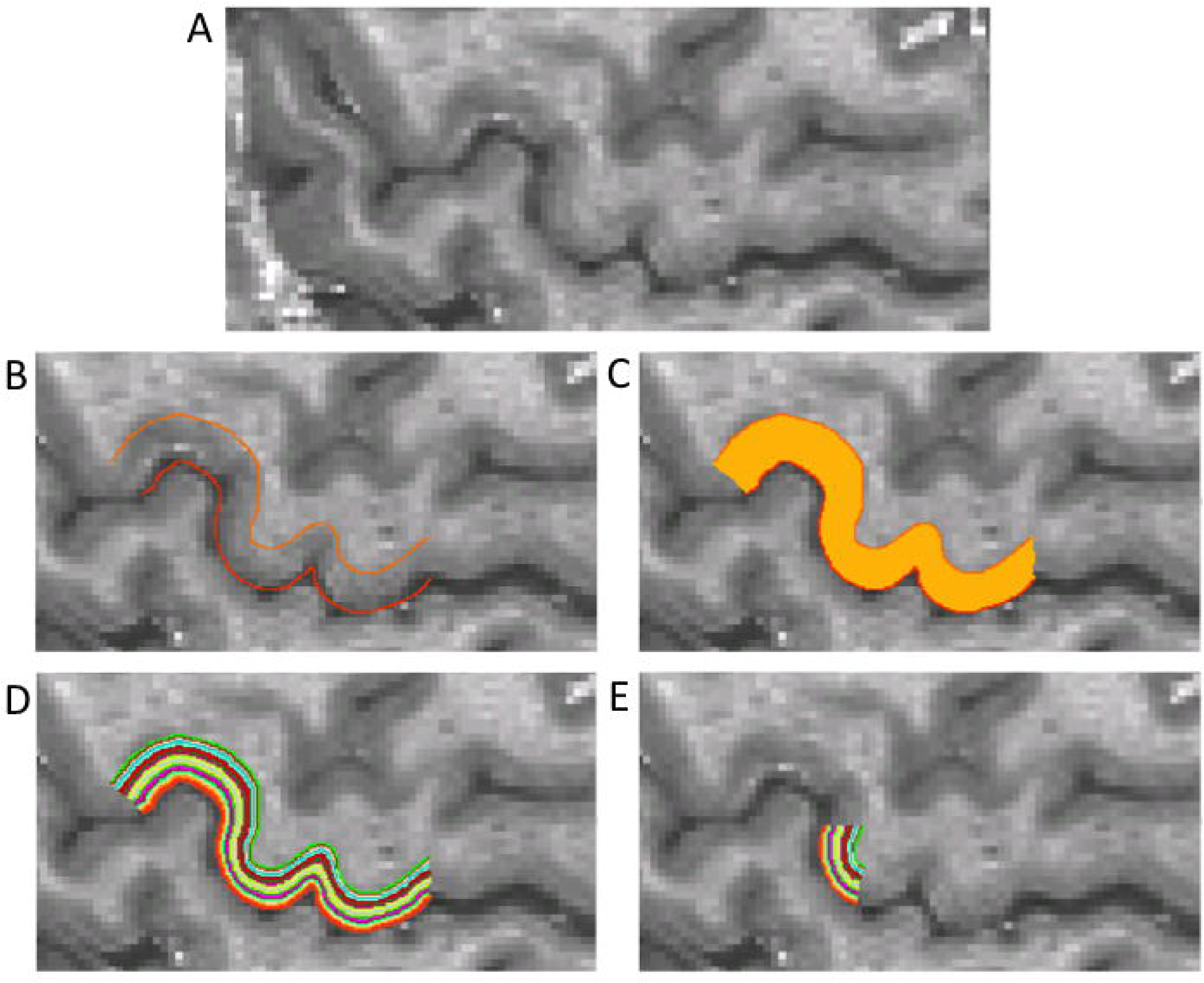
Demonstration of the steps for forming surfaces at different cortical depths directly on the functional images. A) T1-weighted EPI from VASO data. B) GM/WM and GM/CSF boundaries identified on the data. C) Cortical ribbon mask identified. D) Surfaces calculated from GM mask. E) Surface ROIs restricted to lateral bank of M1 hand-knob.

Cortex was divided into 21 equi-distant depths (Fig 4C & D) in the software suite LAYNII and the functional analyses were performed within each, as per previous work (9, 10). Figure 4 shows an example of 21 cortical depths overlaid over T1-weighted functional VASO image. These 21 depths could be used to plot signal change across cortex, as well as to do selective blurring restricted to cortical depth (Fig 6), which aided in visualizing the expected bimodal pattern of activation across depth (10). Activation at different depths was measured in an ROI confined to the lateral side of the hand-knob in M1 (Fig 4E). This is the evolutionary older portion that is located less deep in the central sulcus (aka. ‘old’ M1, rostral M1, or BA4a) (55), which lacks the cortico-motoneuronal cells and is the location of the double peak feature investigated in previous studies (10). Sensitivity and specificity were examined by calculating the average signal change across layers (sensitivity), and by the slope of a line fit to each depth profile that indicated the level of surface bias (an inverse measure of specificity) (10, 56). The effect of sequence type on sensitivity and surface weighting was tested for significance with a repeated measures ANOVA followed by post-hoc analyses. Additional analysis was also done using the FSL FEAT toolbox (57).

**Figure 6:**
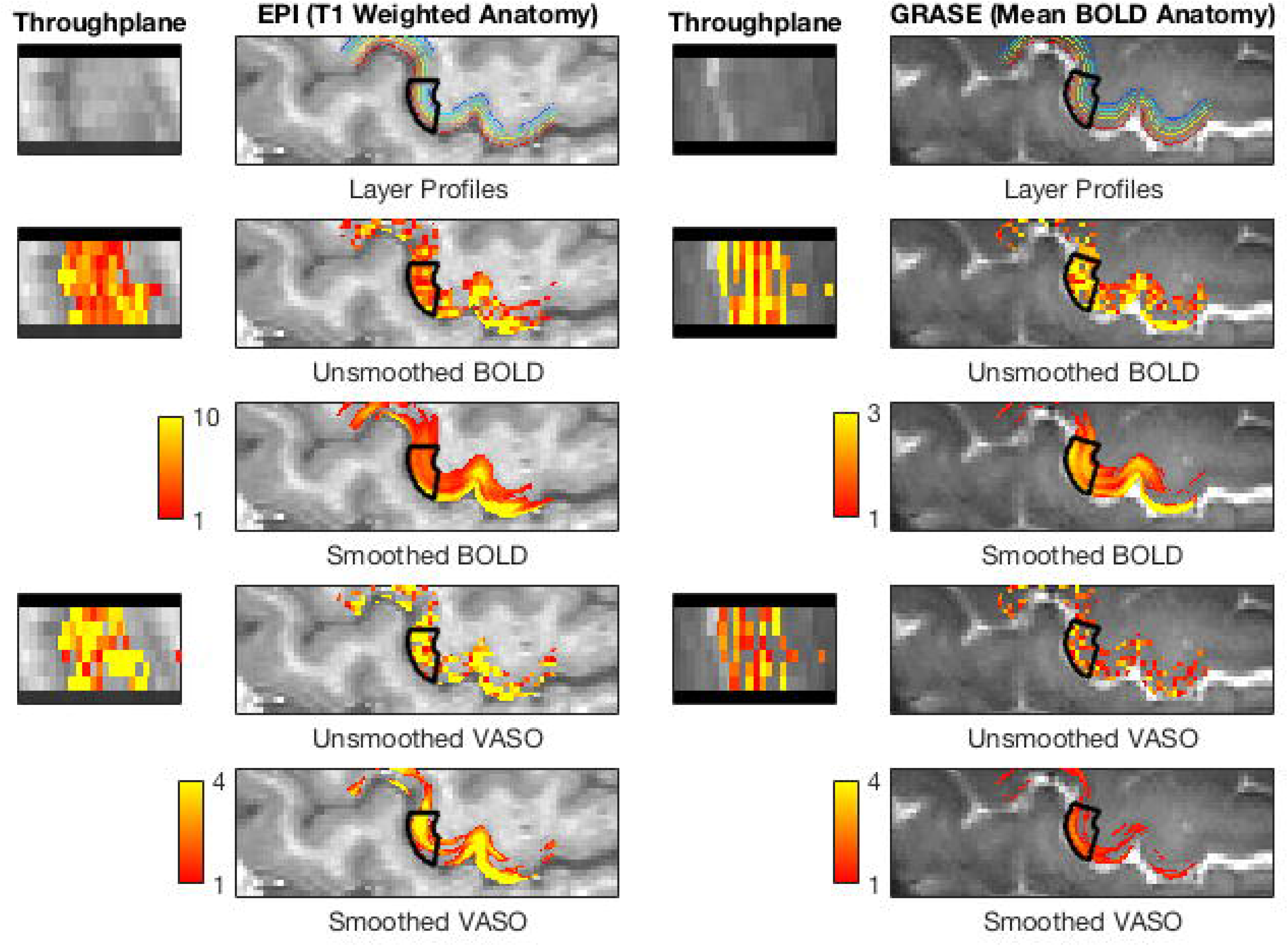
Example signal change maps for an example subject for EPI BOLD/VASO and GRASE BOLD/VASO. Signal change maps are masked by the GM. EPI BOLD maps show highest values around the pial surface, whereas EPI VASO and GRASE BOLD both show two peaks at deep and superficial depths. Note that BOLD and VASO are thresholded differently due to the different contrast mechanisms involved. Black contour indicates the region from which depth profiles were calculated.

#### PSF Simulations

To investigate the signal behaviors of GM tissue in the 3D EPI and 3D GRASE sequences, Bloch equation simulations were performed by numerical application of 3×3 rotation and relaxation matrices, followed by averaging the signal intensity over spin isochromats at each echo time under different regimes of constant flip angle (CFA) and VFA. The simulation parameters were identical to imaging parameters summarized in Table 2 except for the fact that no inversion pulse was applied. The CFA used were 16° for 3D EPI and 165° and 180° for 3D GRASE, respectively. The VFA used for 3D EPI were the same as in the current experiment, and for 3D GRASE were 109°, 63°, 62°, 63°, 75° (calculated by solving an inverse solution of the Bloch equation) (49). Point spread functions for all simulations was numerically estimated by mapping the simulated GM signal evolution into partition direction according to the centric reordering to create a modulation transfer function, and applying the inverse Fourier transformation to the modulation function. The resulting PSFs from the simulation were then compared to those calculated from the acquired data using FSL smoothest (57).

**Table 2).** Imaging parameters for the two sequences used in the current study. Whilst resolution was kept constant, other imaging parameters varied according to the sequence used.

Table 1) Simplifying terms used for the four different image/contrast types.
|  | GRASE | EPI |
| --- | --- | --- |
| FOV (mm) | 99x25x12 | 98x32.8x12 |
| Matrix | 132x34x8 | 132x44x8 |
| Phase Encode | AP | RL |
| In plane resolution (mm<sup>2</sup>) | 0.75 | 0.75 |
| Slice thickness (mm) | 1.5 | 1.5 |
| Slices per slab | 8 | 8 |
| Partial Fourier | 5/8 (Partition) | 6/8 (Phase) |
| GRAPPA | - | 2 |
| TR (between IR pulses, ms) | 3000 | 3000 |
| TR (between image readouts, ms) | 1500 | 1500 |
| TE (ms) | 48 | 24 |
| TI (ms) | 1100 | 1100 |
| Excitation flip angle (degree) | 90 | 26 - 90 |
| Refocusing flip angle (degree) | 165 | - |
| BW (Hz/Px) | 1052 | 1062 |
| Inversion Pulse | BIR4 | TR-FOCI |

## Results

### Task-related signal change

Figure 5 shows the mean relative signal change over 12 block repetitions for 6 volunteers. EPI and GRASE BOLD signal (green and red lines, respectively) increases with activations; EPI and GRASE VASO signal (blue and black lines, respectively) decreases, as an increase in CBV corresponds to a decrease in VASO signal. BOLD EPI shows the largest relative signal change (6.5% +/-1.3), GRASE BOLD and EPI VASO show similar amplitudes (with opposite signs) (3%+/- 0.48, 3%+/-0.43), and GRASE VASO shows the lowest amplitude (2%+/-0.4). Additional data from a separate session where 4 of the subjects were rescanned are shown in Supporting Information Figure S1.

**Figure 5:**
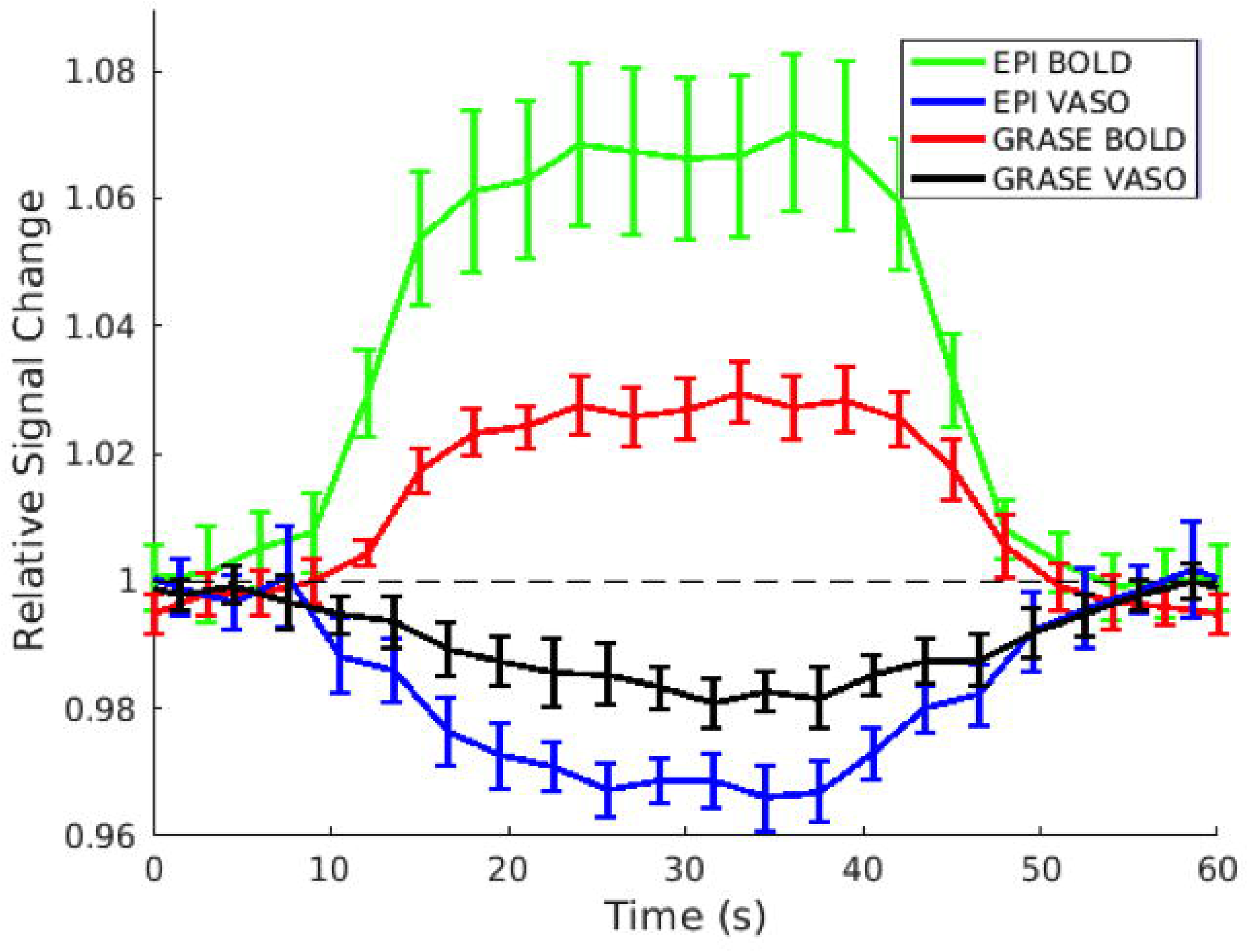
Average timecourse in hand-knob region of M1 across stimulus cycles for EPI BOLD (green), EPI VASO (blue), GRASE BOLD (red) and GRASE VASO (black). BOLD signal change is positive during finger tapping, VASO signal change is negative relating to an increase in CBV. Errorbars = SEM across subjects (N=6).

### Cortical depth functional signal change

Figure 6 shows example signal change maps for EPI and GRASE in a single subject, for both BOLD and VASO contrasts. Unsmoothed and surface-smoothed maps are shown for each contrast/readout comparison. Note that owing to differences in contrast mechanisms between BOLD and VASO, signal change maps are differently thresholded. Additional signal maps for additional subjects are shown in Supporting Information Figure S2. In addition, z-maps from a GLM run on the data using FSL FEAT (57) are included in Supporting Information Figure S3.

Figure 7 shows the average depth profiles for each scan type across subjects, showing differing levels of surface bias and ability to resolve the expected double peak activation profile. Individual depth plots are shown in Supporting Information Figure S4. Each subject shows the expected surface bias for EPI BOLD, and evidence of double peaks for EPI VASO. Double peaks of varying clarity are also seen in the GRASE BOLD and VASO data.

**Figure 7:**
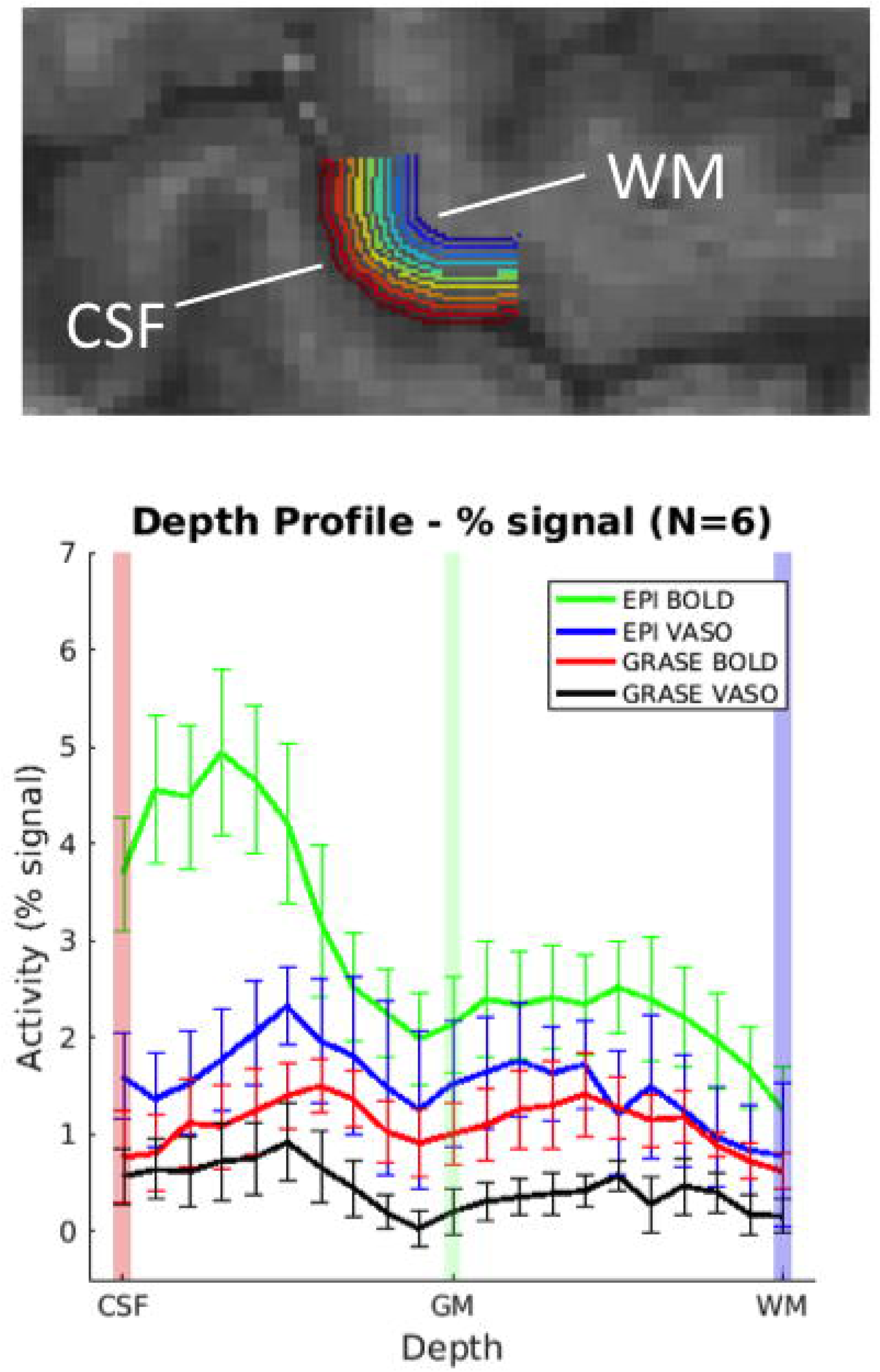
Average depth profiles for signal change between superficial (toward CSF) and deep (towards WM) depths. Color bars on plots indicate signal at depth indicated by corresponding color in example anatomy (top). Errorbars = SEM across subjects (N=6).

Figure 8 compares the sensitivity versus the specificity of the four different scan types. Sensitivity is defined as the mean % signal change across depth for each scan type, and specificity defined as the inverse of slope fit to each profile across depth (10). Figure 8A shows the slopes fit to each mean profile per image contrast type. Figure 8B shows comparisons of sensitivity and surface weighting (inverse specificity) for each contrast. The effect of sequence type on sensitivity and surface weighting was tested for significance with a repeated measures ANOVA with post-hoc analyses. There was a significant effect of sequence type on sensitivity (F_3,15_ = 20.4, p<0.0001) and surface weighting (F_3,15_ = 9.29, p<0.01). Post hoc analyses (Bonferroni corrected) revealed a significant difference in sensitivity between EPI BOLD and GRASE BOLD (p<0.01) and GRASE VASO (p<.01), a significant difference in sensitivity between EPI VASO and GRASE VASO (p<0.05), and a significant difference in surface weighting between EPI BOLD and GRASE BOLD (p<0.05).

**Figure 8:**
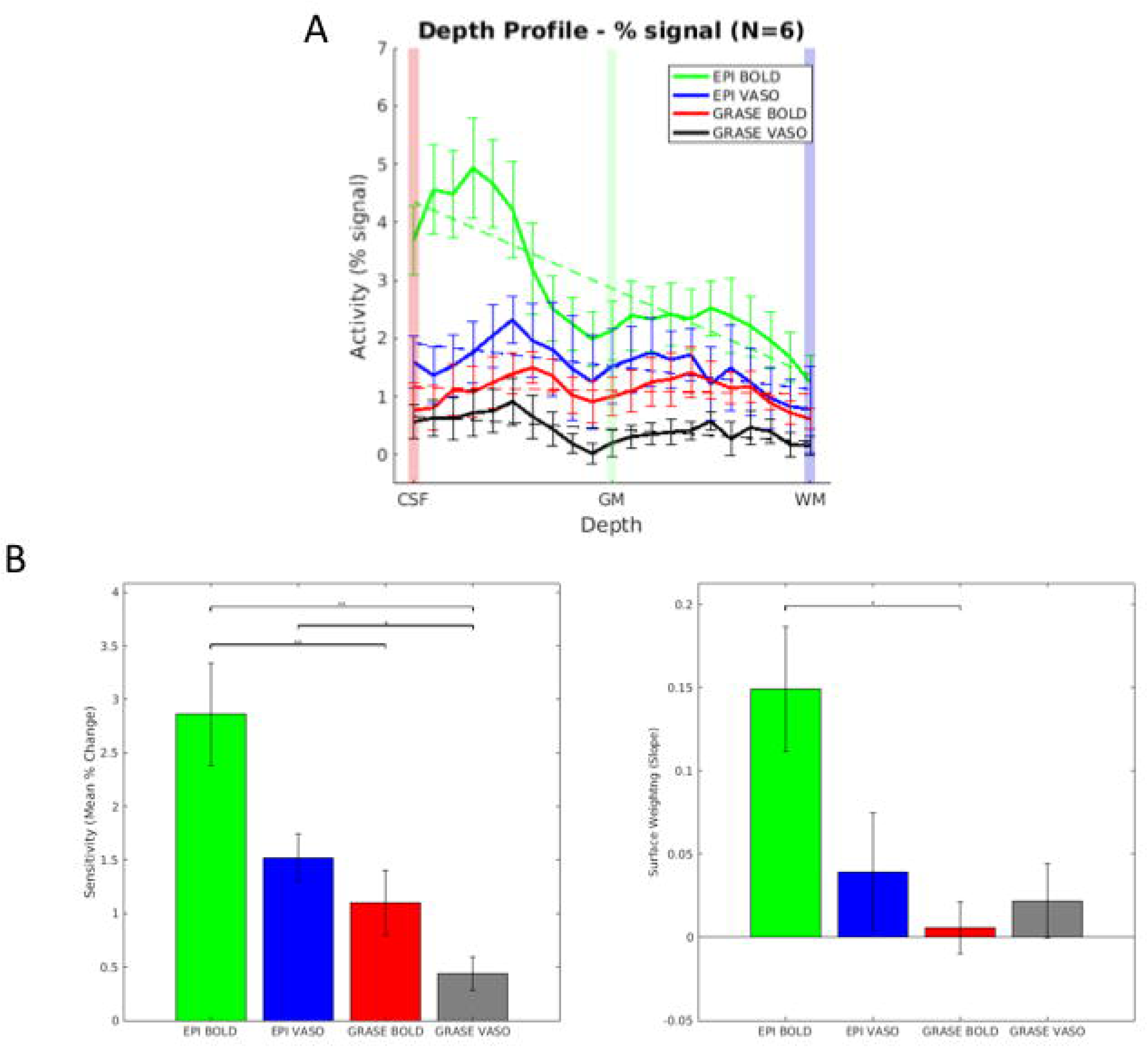
A) Sensitivity and specificity of the four different sequence types. Sensitivity is defined as the mean % signal change across depths, and specificity is defined by the slope a line fit to each profile across depth B) Comparison of sensitivity and surface weighting amongst the four sequences. Post-hoc analysis indicates that EPI BOLD (green) shows significantly greater sensitivity than GRASE BOLD (red) and GRASE VASO (black) and EPI VASO (blue) shows greater sensitivity than GRASE VASO. In addition, EPI BOLD shows significantly greater surface weighting than GRASE BOLD.

Simulations of the signal variation across the echo trains for 3D EPI and 3D GRASE (Fig 9a) yield estimates of the PSF in the slice direction for the two acquisitions (Fig 9b). Due to the signal variation caused by T2 decay in 3D GRASE, the PSF in the slice direction for 3D GRASE is around twice that for 3D EPI, which maintains a flat signal profile across the echo train due to the VFA approach used in this acquisition. FWHM estimates based on FSLs “smoothest” algorithm (57) (Fig 9c) indicate that the FWHM in the partition direction is in fact around two times bigger for the GRASE sequence than the EPI sequence, confirming the estimates from signal simulations.

**Figure 9:**
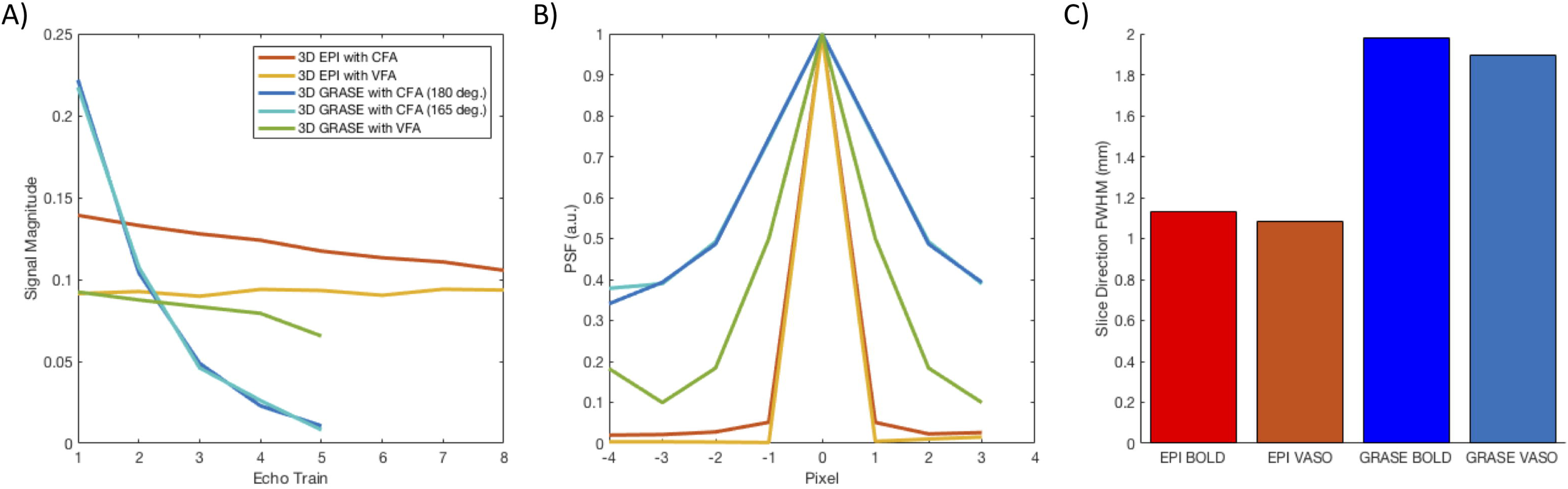
Point Spread Functions (PSF) in the slice/partition direction for EPI and GRASE under different regimes of constant flip angle (CFA) and variable flip angle (VFA) A) Simulated signal across the echo train for 3D EPI without (red) and with (yellow) a variable excitation flip angle component to compensate for T1 recovery, and for 3D GRASE with a CFA (180o, blue; 165 o, teal) and VFA refocusing (green) to compensate for or reduce T2 decay. The use of VFA results in a flatter signal profile across the echo train. B) The resulting PSF from the respective signal decays in echo trains. 3D EPI has a narrower PSF than 3D GRASE with a constant flip angle. The use of variable refocusing flip angle refocusing in GRASE narrows the PSF (green) and would mitigate blurring on the slice axis. C) PSF from the acquired data estimated using FSLs “smoothest” command, confirming the ∼2-fold greater PSF in the slice direction of GRASE BOLD and VASO (blue) compared with EPI BOLD and VASO (red).

## Discussion

Previous studies have demonstrated the utility of CBV-based VASO imaging for high-resolution, layer dependent fMRI, being less weighted towards surface vasculature than GE-BOLD with the latter’s T2*-based contrast mechanism. A similar suppression of signal from surface vasculature has been shown for 3D-GRASE, raising the question of how laminar specificity compares between VASO and GRASE. We extend previous work combining GRASE and VASO at standard resolutions to UHF, sub-millimeter imaging for laminar analysis.

Comparing relative signal changes in M1 for each contrast/readout (Figure 5), standard EPI-BOLD showed the highest levels of signal change, on the order of 6%, with GRASE-BOLD showing lower levels of around 3%, in line with previous measurements (21, 58). The levels of signal change for EPI-VASO were ∼3%, similar to previously seen changes using EPI (9, 10). Signal change in GRASE-VASO was lowest at around 2%.

The lower signal change for GRASE BOLD versus EPI BOLD is expected given that EPI BOLD contains susceptibility contrast changes from both large and small vessels, whereas GRASE primarily has signal change from only small vessels. In addition, GM signal (and hence VASO contrast) is maximized with the VASO acquisition used here when the overall TR is kept minimal. In this case the time between image acquisition was 1.5s, which was insufficient for full recovery of the SE GRASE signal in GM using full 90 RF excitation and may have led to suboptimal BOLD imaging using GRASE. Without the need for a VASO acquisition component, longer TRs of 2 to 3s seconds (increasing the delay between BOLD readouts, with the same overall number of samples for BOLD imaging as in the VASO acquisition) would allow greater signal recovery for GRASE BOLD imaging, and have been utilized at 7T in mesoscale fMRI studies (5,6,21,33). This lack of overall image SNR may have also led to the lower VASO contrast seen using GRASE-VASO compared to EPI-VASO, despite equivalent levels of signal change observed in previous implementations (39). In theory a single TI, defined by the single excitation pulse for each single-shot 3D GRASE readout, would have led to more consistent blood nulling across k-space, and hence greater quantification of CBV (after BOLD correction using the second image readout) in the GRASE VASO signal compared to 3D EPI, in which the multiple excitations for k-space partitioning lead to inversion times that are not at the blood nulling time for some parts of k-space. However, the lower signal in GRASE-VASO obscured these smaller effects of imperfect blood nulling compared to segmented 3D EPI. The timing parameters (TI/TR) for GRASE VASO were based on those previously optimized for EPI VASO (10), and it may be the case that a different implementation could lead to similar amplitudes for EPI and GRASE VASO, as seen previously at 3T. The short TR and subsequent necessity of a reduced FA for refocusing pulses also reduced the overall SNR for both GRASE BOLD and GRASE VASO imaging, and optimized parameters for GRASE BOLD imaging sequence alone, without being incorporated into VASO acquisition, would further improve layer specificity.

Another reason for the difference in relative signal response of 3D-EPI VASO and 3D-GRASE VASO might be from a different signal evolution along the finite readout duration. The 3D-EPI VASO readout consists of multiple excitations along the inversion-recovery relaxation. While the k-space center is acquired at the TI of the blood nulling time, outer k-space segments can contain residual blood signal. Dependent on the water permeability between intravascular and extravascular space, residual blood signal can accumulate in the extravascular space during the readout and, thus, amplify the VASO contrast. This extra-vascular perfusion component is to some extent balanced out by exchange in the other direction, but simulations show that the net result is an amplification of CBV estimates (7, 59). This perfusion-dependent signal weighting is not to be confused with potential CBF-dependent inflow effects of fresh, not-inverted blood (described in (60)). Such inflow would reduce the VASO contrast, while the permeability effect here refers to perfusion related CBV amplification. While this effect across the short readout train used for 3D EPI should be minimal, the fact that GRASE VASO uses only one excitation pulse means that its T1 signal represents a true snapshot along the inversion-recovery relaxation. Thus, in theory, it should be a more quantitative CBV contrast, without such permeability-related amplifications.

It is also worth noting that based on the different spin-history in 3D-EPI VASO and 3D-GRASE VASO, they are differently sensitive to potential contamination of dynamic CSF volume changes (61, 62). Here, we do not believe that such hypothetical CSF contaminations can explain the different signal changes in our results. First, we previously found that for local activation in the motor cortex during finger tapping, the CSF volume change is negligibly small compared to global systemic respiration tasks (9). Second, such a CSF contamination should be isolated to partial voluming in superficial layers. The profiles in Figure 7, however, show that the 3D-GRASE VASO has a reduced signal change across all cortical depths. Thus, CSF contaminations can be ruled out.

Maps of signal changes for the four sequence types in an example subject (Figure 6) and the average profiles of signal change across depth for the four contrasts (Figure 7) mirror the pattern seen in the time course (Figure 5) results. EPI BOLD has overall higher signal change, but the larger signal change is biased towards superficial cortical depths, as seen previously in humans (9,10,21,22). This indicates that EPI BOLD is mostly influenced by the signal change in large draining veins at the pial surface, non-specific to the location of neural activity. EPI VASO shows little to no surface bias and shows peaks at both superficial and deep cortical depths, as seen in previous studies (9, 10). Similarly, GRASE BOLD does not show a superficial bias, in line with previous studies (4, 21), and also shows the double peak pattern (Figure 7) visible in EPI VASO. GRASE VASO shows the same pattern to some extent although with higher levels of noise, mirroring the lower CNR seen in the time course plots (Figure 5).

The double peaks observed in the BOLD GRASE data are of lower height than those observed with EPI VASO (Figure 7). The depth profiles in both GRASE BOLD and VASO for individual subjects (Supporting Information Figure S4) demonstrate overall larger variations in amplitude compared with those using EPI VASO. In addition to the suboptimal SNR arising from the timing parameters used, this may be due to the broader point spread function (PSF) in the slice direction for 3D GRASE, and dependent larger partial volume effects in cases where the slice placement is suboptimal, i.e. not completely perpendicular to the cortical hand knob region of M1. A second issue is that cortical surfaces were defined separately for the EPI and GRASE scans in each subject to analyze data in native functional space, and in some cases the GM/WM contrast was less clearly defined for GRASE data. This could have led to inconsistencies in placing the GM/WM boundary when defining surfaces, making the peaks in the group average profile less distinct even when seen in individual depth profiles.

We compared sensitivity (mean signal change across depths) and specificity (slope of a linear fit to the signal change across depth) (10) for each contrast type (Figure 8A), and found a significant effect of sequence type on both sensitivity and specificity, with post-hoc analyses revealing EPI BOLD to have a higher sensitivity than other sequences, but also a higher surface weighting (i.e. less specificity). It should be noted that the definition of sensitivity and specificity used here are not entirely separable, as the case of absolutely no signal would be treated as highly specific (i.e. a flat line), so for this analysis sensitivity and specificity should only be considered together. This definition of specificity has been used previously in studies examining laminar responses using fMRI (10,21,56), and a previous comparison with alternative measures of specificity found unchanged results (56).

While it should be noted that key imaging parameters (acceleration, zooming, phase encode direction etc.) differ across the two imaging readouts used (EPI and GRASE), each set of parameters were optimized for the sequence in question to obtain high in-plane resolution for laminar imaging. The EPI sequence parameters were optimized based on a previous experiment in M1 (10), and these parameters would not be optimal for a GRASE sequence, so GRASE was optimized separately (within the constraints of a VASO sequence). Thus, the experiment was not to compare two sequences identical except for their contrast mechanisms, but to compare two sequences separately optimized for mesoscale imaging.

Previous comparisons of VASO and SE-BOLD showed the latter still having a residual surface bias when compared to VASO, albeit less than that seen in GE-BOLD (10). This seems contradictory to the lack of surface bias seen using a SE sequence (GRASE) in the current study, however the difference likely arises from the use of much shorter EPI echo trains in the zoomed GRASE sequence than SE-EPI in the previous comparison. SE-EPI is known to have large vessel contributions arising from T2* contamination in the extended echo train at high resolutions (12). The use of inner volume zoomed GRASE has been shown to decrease surface bias when compared with non-zoomed SE-EPI (4), and the increased specificity seen with GRASE compared with that previously demonstrated with SE-EPI replicates this. Another difference between SE-EPI and GRASE is the T1 contrast contribution from STE in GRASE, not present in single refocusing SE-EPI, which increases with reduced flip angle. In our experiments a nominal flip angle of 165° instead of 180° refocusing pulse was necessitated to stay within SAR limits using the 1.5s effective TR between GRASE readouts.

VASO is also not completely independent of macrovascular biases towards the pial surface. As previously discussed in (Figure 8 of (9),(63)), the larger vascular density of diving arterioles and micro-vessels in superficial and middle layers can result in higher signal changes compared to deeper layers.

The results of this work show that spatial specificity of both GRASE-VASO and GRASE-BOLD is sufficient for laminar specific imaging, yielding two distinct functional activation peaks across the depth cortex in M1 (Figure 7), in agreement with underlying input and output-driven activity associated with a finger-tapping task.

The use of different image contrasts for laminar fMRI in humans has been of increasing interest since mesoscale resolution became achievable. The weighting towards pial vessels shown for GE EPI makes the straightforward resolving of BOLD signals from specific laminae difficult (21, 22), necessitating that superficial depths, non-specific voxels, or voxels containing veins be excluded during analysis (23), or necessitating unique experimental design (64, 65) or analysis (5, 58) to begin to resolve layer specific signals. The use of SE contrast at high field strengths to suppress signals from large veins has been combined with 3D imaging in GRASE (2) to demonstrate mesoscale fMRI without this bias towards the pial surface (4, 21). GRASE has been used to demonstrate consistent selectivity for a given stimulus property across depth (the hallmark of cortical columns) in various cortical areas (6,34,58), and to demonstrate certain cortical computations restricted to certain cortical depths (5, 33). The current study is the first to show two distinct activity peaks in human M1 in BOLD fMRI, which has been used as a hallmark of laminar selectivity when assessing different contrasts for laminar fMRI (10).

Previous work suggests that VASO (8) acquisition methods for fMRI have an optimal tradeoff of functional sensitivity and spatial specificity (7, 10). However, VASO imaging requires implementing an adiabatic inversion recovery RF pulse, which has to be capable of providing inversion with decreased inversion efficiency [e.g. a BIR4 (51) or TR-FOCI (66) pulse] and increases the acquisition time by blood nulling time, which is in the order of 1 s. It also requires a very careful timing implementation, so that no fresh (non-inverted) blood flows into the imaging volume, which requires the arrival time and T1 of blood in a given brain area to be known. While variable blood T1 across subjects has been reported, the effects of these small variations in T1 are unlikely to have a large effect on the VASO signal (9). In addition, there is residual BOLD contrast in the blood-nulled images, especially at ultra-high-fields, which is of the opposite sign and frequently of higher amplitude than the negative VASO signal. This problem can be alleviated using the BOLD-corrected VASO (7) method, where two images with and without blood-nulling are required, and the latter, purely BOLD weighted image is used to correct the residual BOLD signal in the blood-nulled image. However, this requires additional image post-processing and creates the problem of reduced temporal resolution of the VASO acquisition as compared to the BOLD images alone. Further development and sequence parameter optimization of GRASE alone could have advantages to VASO as it would eliminate the use of complex sequence timing with inversion pulses and the inefficiency of acquiring a double readout for BOLD T2* correction. A recent development of compressed sensing (CS) 3D GRASE to increase slice coverage is a promising further development of the methods used here (67).

Although the sensitivity and specificity of GRASE demonstrated here make it a good candidate for mesoscopic imaging, it is not without drawbacks. As noted above, the PSF in the slice direction is increased for GRASE (Figure 9) due to the T2 decay across the echo train which can be mitigated to an extent using VFA on the refocusing pulses (49), at the expense of some overall SNR. These differences in PSF between the used sequences could affect the sensitivity and specificity as measured in this paper, for example through partial volume effects. However, in this particular experiment, the larger PSF in the slice direction for GRASE will have less of an effect than it might otherwise owing to the nature of the neural anatomy under investigation, where thicker slices were placed orthogonal to the central sulcus and laminar activity was resolved inplane. In more convoluted areas of cortex where resolutions closer to isotropic are required, methods to mitigate through plane blurring in GRASE such as VFA (49) or CS (67) may prove useful.

The question still arises as to whether these T1 contributions from STE in GRASE increase the risk of inflow artifacts (68, 69). Although on the one hand the longer T1 of blood at high fields (50) may lead to an increased chance of inflow artifacts, the simultaneous shortening of blood T2/T2* is likely to counteract this. In addition, as GRASE is a 3D sequence, with orthogonal slab excitation and refocusing volumes, this should also reduce the potential impact of inflow artifacts. Additionally, only short (5 refocusing pulses) SE trains were used in these GRASE experiments, giving less opportunity for STE magnetization to build up in the later echo train.

In addition to the inflow effects commonly seen in BOLD images, all VASO sequences are at risk of artifacts due to the inflowing of uninverted blood to the imaging volume, e.g. in areas of very short arrival time such as large arteries (70), potentially reducing any observed VASO signal change. The use of shorter TIs with adjusted inversion efficiency, as performed here, can help avoid these artifacts.

### Conclusion

The results presented here demonstrate the feasibility using a 3D GRASE readout at ultra-high field for cortical layer fMRI at the mesoscale. The sensitivity and specificity in laminar fMRI achieved with BOLD using 3D GRASE is similar to that in CBV-based imaging using VASO with 3D GE-EPI readout. 3D GRASE BOLD uses inner volume zooming which also reduces the T2* decay in the signal acquisition to largely remove the contribution of pial draining veins, providing laminar specificity with BOLD contrast.

The ability to specifically measure neuronal signal in cortical layers provides a unique approach to studying circuitry in human brain where different cortical layers act as inputs from, or send outputs to other cortical areas. By mapping this mesoscale organization in brain circuitry, cortical layer fMRI can be used to study the brain’s directional hierarchy in long range circuits between the hundreds of distinct brain areas and the microcircuits within cortical layers. Therefore, cortical layer fMRI may also play an essential role in understanding the dysfunction of brain circuits affecting higher-order brain functions.

## Acknowledgements

NIH BRAIN Initiative grants - 5U01EB025162, 1R24MH106096, 5R01MH111444, 1R01MH111419, 5R44NS084788. Laurentius Huber was funded form the NWO VENI project 016.Veni.198.032 for part of the study. We thank Matthias Gunther for his advice on rf pulses.

## Supporting Information

Supporting Information Figure S1 – Percent signal change (left) for the four sequences, also shown scaled to the maximum of the highest curve (right) for the data in the main study (top) and a follow up session in 4 subjects (bottom).

Supporting Information Figure S2 – Smoothed and raw signal change maps for individual subjects for EPI BOLD/VASO and GRASE BOLD/VASO. Note that BOLD and VASO are thresholded differently due to the different contrast mechanisms involved.

Supporting Information Figure S3 – Smoothed and raw Z-Score maps for individual subjects for EPI BOLD/VASO and GRASE BOLD/VASO.

Supporting Information Figure S4 – Depth profiles for individual subjects for EPI (top row) and GRASE (bottom row) for BOLD and VASO.

