## Supplemental Figures for "Comparison of BOLD and CBV using 3D EPI and 3D GRASE for cortical layer fMRI at 7T"

Session 1 (6 Subjects)

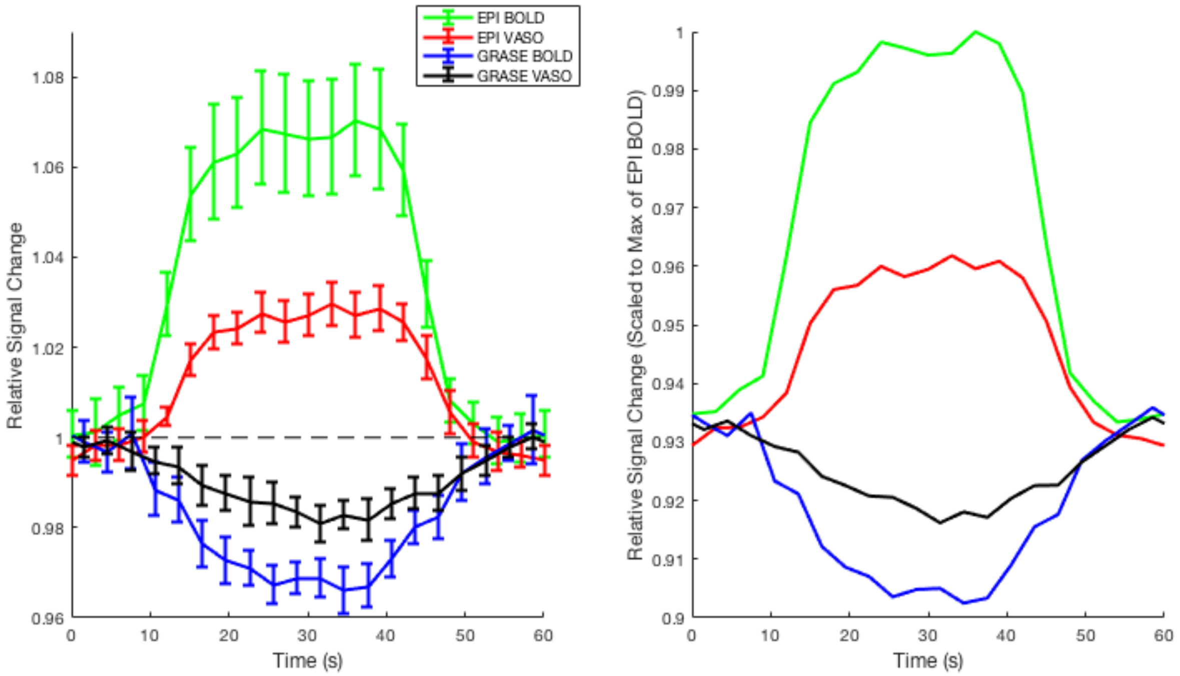


Session 2 (4 Subjects)

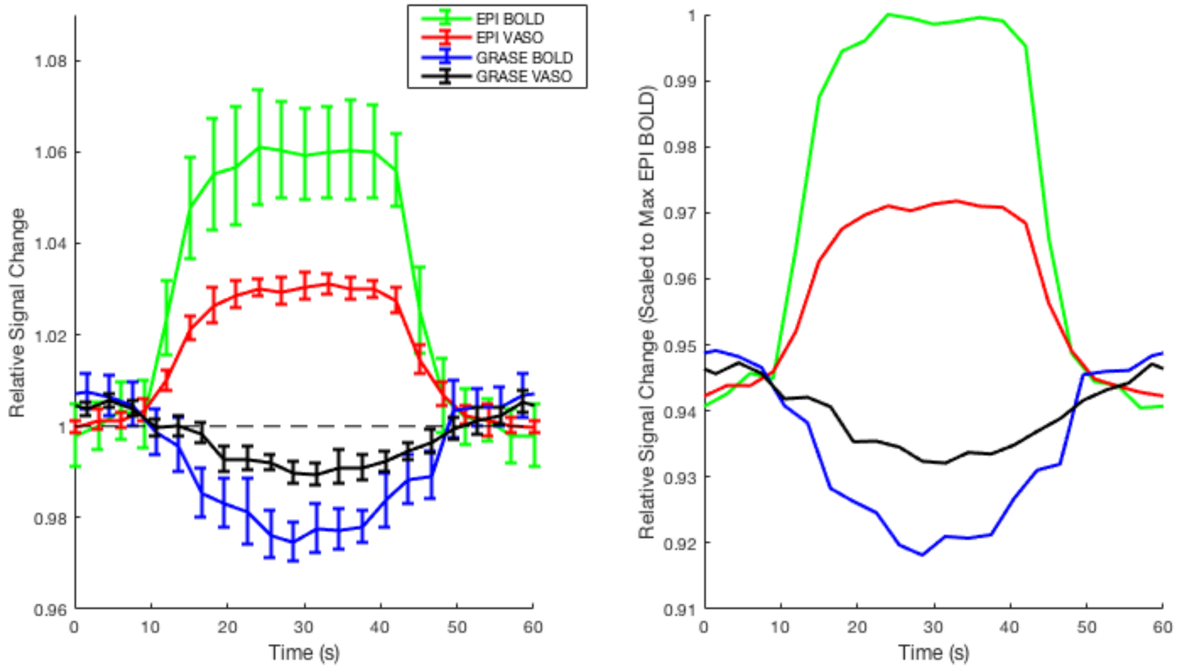


Supporting Information Figure S1 – Percent signal change (left) for the four sequences, also shown scaled to the maximum of the highest curve (right) for the data in the main study (top) and a follow up session in 4 subjects (bottom).

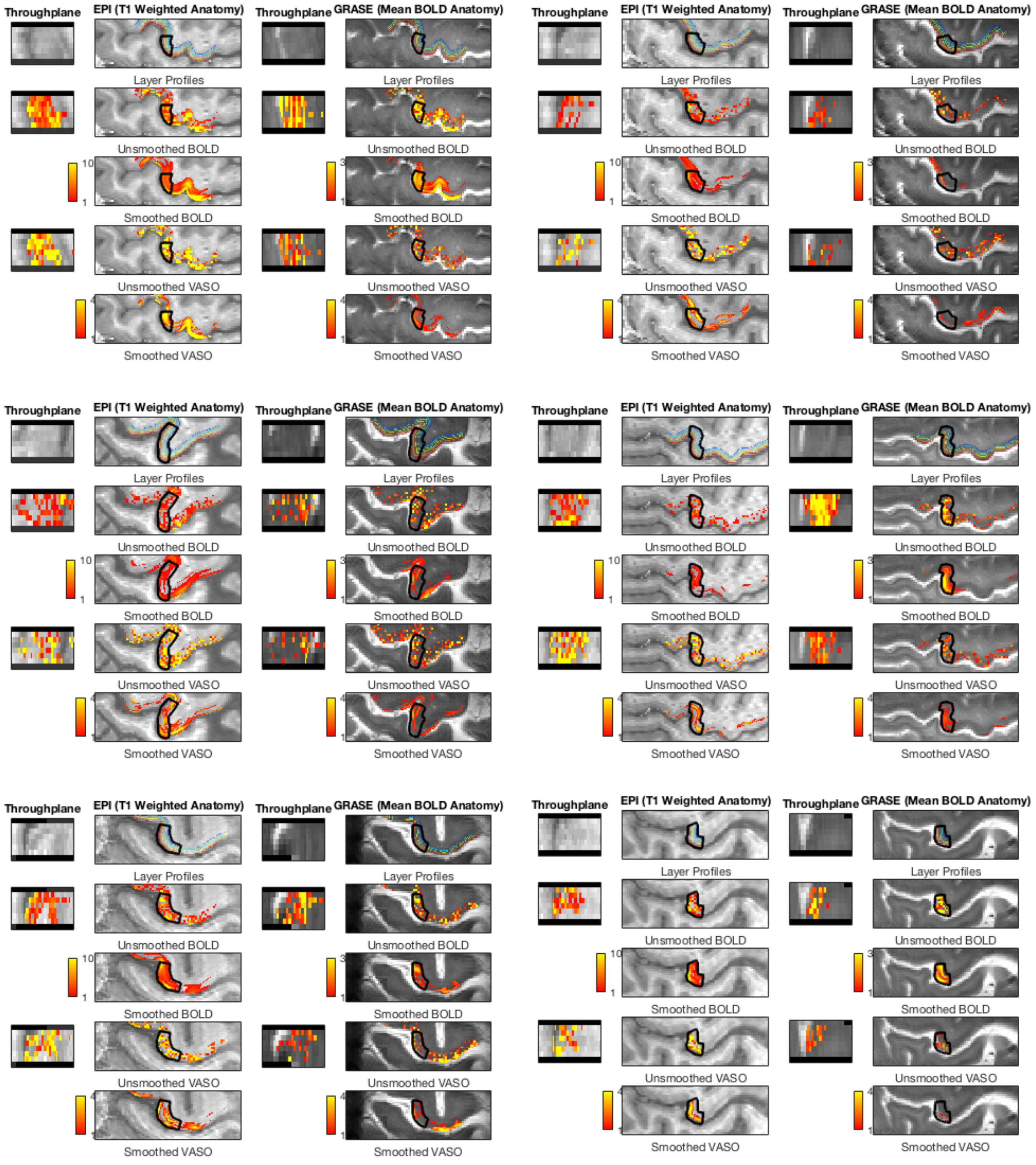


Supporting Information Figure S2 – Smoothed and raw signal change maps for individual subjects for EPI BOLD/VASO and GRASE BOLD/VASO. Note that BOLD and VASO are thresholded differently due to the different contrast mechanisms involved.

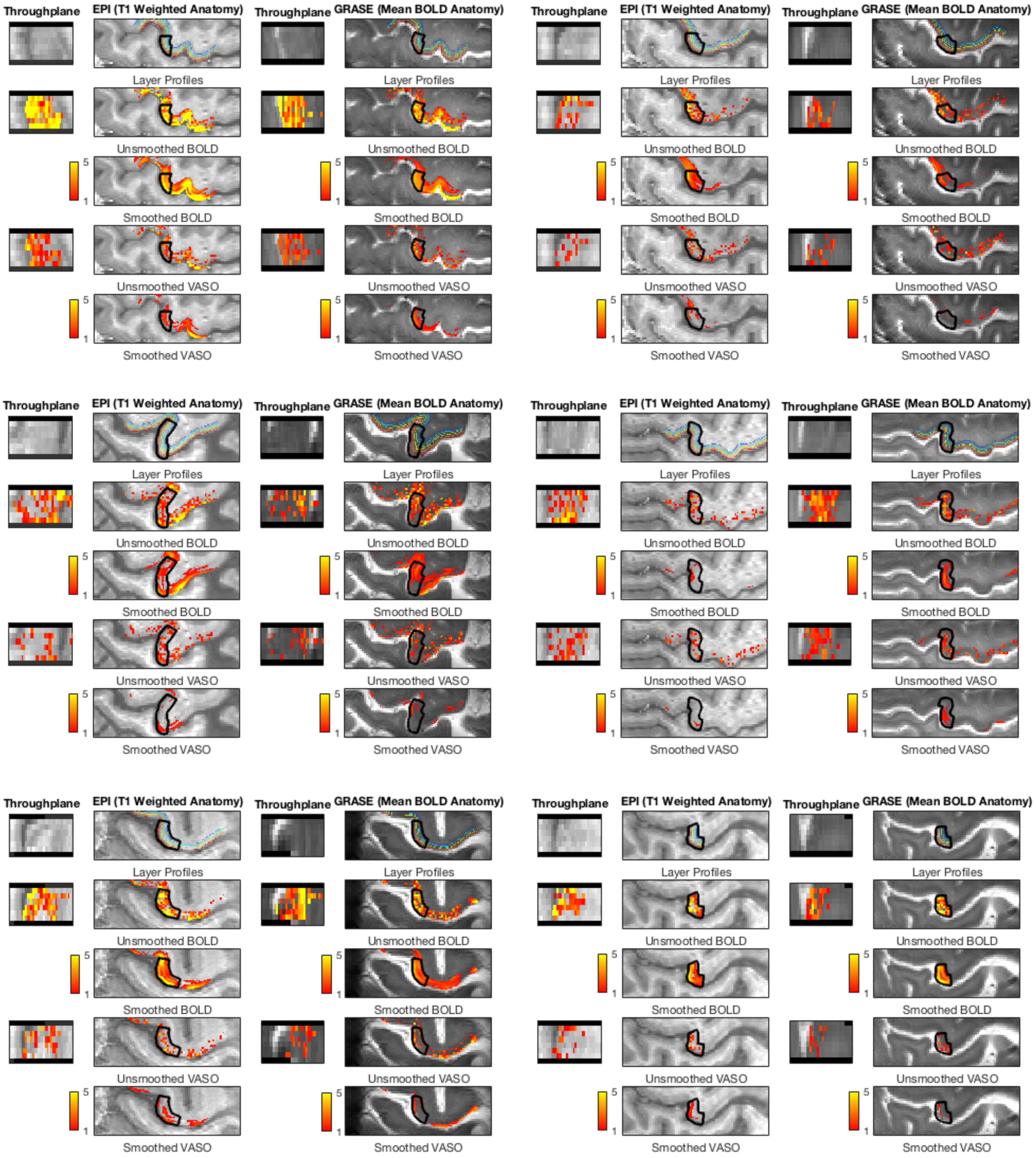


Supporting Information Figure S3 – Smoothed and raw Z-Score maps for individual subjects for EPI BOLD/VASO and GRASE BOLD/VASO.

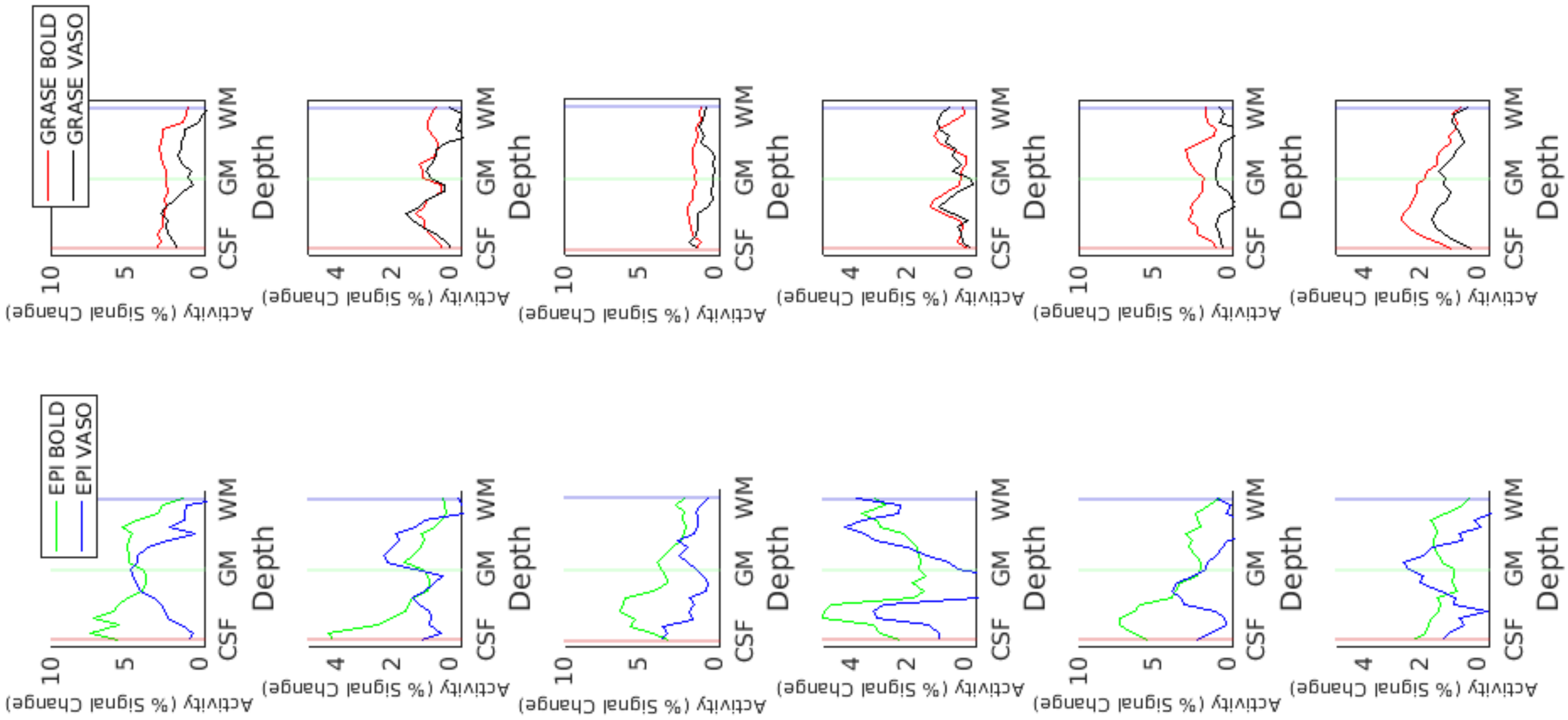


Supporting Information Figure S4 – Depth profiles for individual subjects for EPI (top row) and GRASE (bottom row) for BOLD and VASO.
